# The cognitive basis of intracranial self-stimulation of midbrain dopamine neurons

**DOI:** 10.1101/2022.08.11.503670

**Authors:** Samuel J. Millard, Ivy B. Hoang, Zara Greer, Shayna L. O’Connor, Kate M. Wassum, Morgan H. James, David J. Barker, Melissa J. Sharpe

## Abstract

Recently there has been a reckoning in the dopamine field. This has suggested that the dopamine prediction error may function as a teaching signal, without endowing preceding events with value. We studied the cognitive basis of intracranial self-stimulation (ICSS), a setting where dopamine appears to be valuable. Physiological frequencies seen during reinforcement learning did not support robust ICSS or promote behavior that would indicate the stimulation was represented as a meaningful reward in a specific or general sense. This was despite demonstrating that this same physiologically-relevant signal could function as a teaching signal. However, supraphysiological frequencies supported robust ICSS where the stimulation was represented as a specific sensory event, which acted as a goal to motivate behavior. This demonstrates that dopamine neurons only support ICSS at supraphysiological frequencies, and in a manner that does not reflect our subjective experience with endogenous firing of dopamine neurons during reinforcement learning.

**One sentence summary:** Dopamine neurons only support ICSS at supraphysiological frequencies and in a manner not reflecting dopamine’s role in learning.

## Main Text

For many decades intracranial self-stimulation (ICSS) has been used to investigate the neural substrates of motivation^1–9^. The finding that many species, including humans, will work to receive stimulation of various brain regions has been taken as evidence that those regions encode natural rewards or pleasure^5,6,8,10^. Regions that support ICSS predominantly rest across the medial forebrain bundle, but can also be found in many parts of the limbic and extrapyramidal motor systems^1,10–13^. The use of this paradigm for studying the neural substrates of reward has increased considerably in the last decade, afforded in part by the optogenetic revolution that allows for incredibly precise control over neuronal activity in rodents^14–18^. This has extended our knowledge of the particular neuronal populations responsible for the ICSS effect, down to the specific projection targets that subserve these behaviors^12,13,15,19–25^.

One of the primary neuronal populations that promote ICSS are midbrain dopamine neurons. Rodents, primates, and even humans will work vigorously to receive stimulation of dopamine neurons^3,4,7,23,25–28^. This has been taken as strong evidence in favor of the value hypothesis of dopamine, which argues that phasic dopamine assigns scalar value to external stimuli or actions antecedent to dopamine firing to encourage motivated behavior directed towards rewards^29–34^. The value hypothesis of dopamine has been a long-standing dogma in the field. However, recent findings have suggested that this is not the whole story. Specifically, many published results now show that dopamine transients act as teaching signals to help us form associative maps of how stimuli relate to one another^22,35–44^, without making those stimuli valuable^36^. These data call into question the idea that phasic dopamine activity is solely dedicated to value assignment and require a more nuanced model, which is an endeavor reflected by several recent efforts in the field^45,46^.

Apparent differences in the function of phasic dopamine may also result from the way in which dopamine neuronal activity is manipulated in different settings. For example, during ICSS, stimulation of dopamine neurons is delivered when the subject has completed an action. Yet the brief increases in the phasic firing of dopamine neurons that characterize the prediction error are generally seen at the unexpected transition between two events (e.g. cues and rewards, or actions and rewards) ^29,47–50^, and not at the completion of an inconsequential action. Effectively, the way a prediction error acts when couched in the context of learning to associate two things together in the physiological world is likely different to how it might function to produce ICSS^36^. The second is that the quintessential dopamine prediction error signal, famously revealed by Schultz and colleagues in the nineties^29^, is very brief and typically observed to be within the range of 10-20Hz in primates and rodents^29,47,51^. However, ICSS stimulation parameters are generally much higher than the range seen during reinforcement learning; conventionally, dopamine stimulation with ICSS is around 50Hz^4,7,52,53^, and studies often use frequencies above even this supraphysiological range^2,4,28,52^.

To begin to understand why phasic activity of dopamine neurons subserve different functions in ICSS relative to reinforcement learning, we tested the cognitive basis of ICSS of dopamine neurons using both learning-relevant and supraphysiological frequencies of dopamine stimulation. This would allow us to simultaneously ask whether dopamine stimulation acts as a general value signal to produce ICSS, consistent with the value hypothesis^29^, and if the role of dopamine in ICSS varies with stimulation parameters. Surprisingly, despite the thousands of manuscripts that have demonstrated ICSS, there are very few findings that have investigated the question of *how* this stimulation is represented in the brain. To test the nature of the representation supporting ICSS, we investigated how dopamine stimulation would interact with the well-studied Pavlovian-to-Instrumental transfer (PIT) effect, which is seen in both rodents and humans, and allows for a differentiation between specific and general representations of rewards^54–58^. Specifically, PIT allows us to determine whether a reward is supporting learning because it acts as an internal representation of a goal, or whether it simply reinforces a response via a more general valence mechanism. The PIT procedure involves first teaching subjects that two cues lead to different types of rewards (e.g., grain pellets or sucrose). Then subjects learn separately that they can work to earn these two rewards by performing two different actions (e.g., a left or right lever press). Finally, subjects are given a test where the cues are played and two actions are available but do not produce their associated reward. This test is what allows us to probe the nature of the reward representation that contributed to learning in the earlier phases of the task. Specifically, if the reward was capable of driving learning by evoking a sensory-specific representation of itself, then presenting the cue should motivate the subject to perform the action associated with the same reward. For example, if the pellet-predictive cue is presented, it will make the subject think of the pellet, and motivate them to press the lever that is also associated with pellets. However, if learning was devoid of a specific representation of reward identity, and driven solely by a more general value mechanism, cue presentation will motivate lever-press responding in a non-specific way, increasing responding on both levers^56,58^. Generally, food rewards will support selective PIT, indicating the presence of a specific association over and above a more general value mechanism^56,58^. Armed with this knowledge, we redesigned this selective PIT design in rats to include dopamine stimulation as one of the rewards and compared this with a food reward. Given the value hypothesis argues that the dopamine error signal seen to unexpected food rewards reflects the magnitude of the reward’s scalar value^29^, this would allow us to directly compare how the signal functions as a reward in its own right relative to the reward it accompanies. We included two groups, one that received dopamine stimulation approaching physiological levels during reinforcement learning (20Hz)^36,37,51^, and another at supraphysiological rates that are typically used to show ICSS (50Hz)^4,7^. This allowed us to test how if dopamine stimulation is represented as a specific reward within the brain, how it compares to a natural food reward, and whether this differs for physiological and supraphysiological frequencies.

Prior to training, we infused 2ul of a Cre-dependent excitatory opsin channelrhodopsin (ChR2; pAAV5-Ef1a-DIO-hChR2(E123T/T159C)-eYFP) bilaterally into the ventral tegmental area (VTA) of male and female rats expressing Cre-recombinase under the control of the tyrosine hydroxylase (TH) promoter^36,37^ (*n*=11; Fig 1A-C; for power analyses see methods and Fig S1). During this surgery, we also implanted optic fibers bilaterally into the VTA. This allowed us to stimulate activity in dopamine neurons in the VTA via blue light delivery (473nm,16mW, 1s)^16^. Four weeks after surgery, rats were placed on mild food restriction. Then, training began with presentations of two 2-min auditory cues (white noise or click) on separate trials, intermixed within a session, across 10 days. During presentation of these cues, the associated reward was delivered 4 times, randomly distributed through cue presentation. All rats received one cue paired with dopamine stimulation (20Hz or 50Hz) and the other with sucrose pellets, counterbalanced. We recorded locomotor activity as well as the number of entries rats made into the food port during cue presentation (Fig 1G and J). Locomotor activity increased across sessions for both the dopamine and food-paired cues (main effect, session: *F*_4,36_=8.595, *p*<0.0001), with no difference in the degree of locomotor activity evoked by the cues (main effect, dopamine vs. food cues: *F*_1,9_=0.075, *p*=0.790), or any between-group difference in locomotor activity (main effect, 20Hz vs. 50Hz groups: *F*_1,9_=0.482, *p*=0.505; Fig 2A and D). Unsurprisingly, entries into the food port were only seen for the food-paired cue and not the dopamine-paired cue. A repeated-measures ANOVA on the food-port-entry data showed a significant main effect of cue (dopamine vs. food: *F*_1,9_=13.221, *p*=0.005), with no interaction by group (*F*_1,9_=0.105, *p*=0.753). Similarly, there was a main effect of session (*F*_4,36_=10.425, *p*=0.000), and a session by cue interaction (*F*_4,36_=9.196, *p*=0.000) with no interactions by group (session x group: *F*_4,36_=0.522, *p*=0.720; session x cue x group: *F*_4,36_=0.262, *p*=0.900). Finally, there was no between-group difference in overall responding in the food port to the cues (*F*_1,9_=0.106, *p*=0.752). This shows that locomotor activity increased across learning to both cues in both groups, and only the food-paired cue promoted entries into the food port.

**Figure 1.**
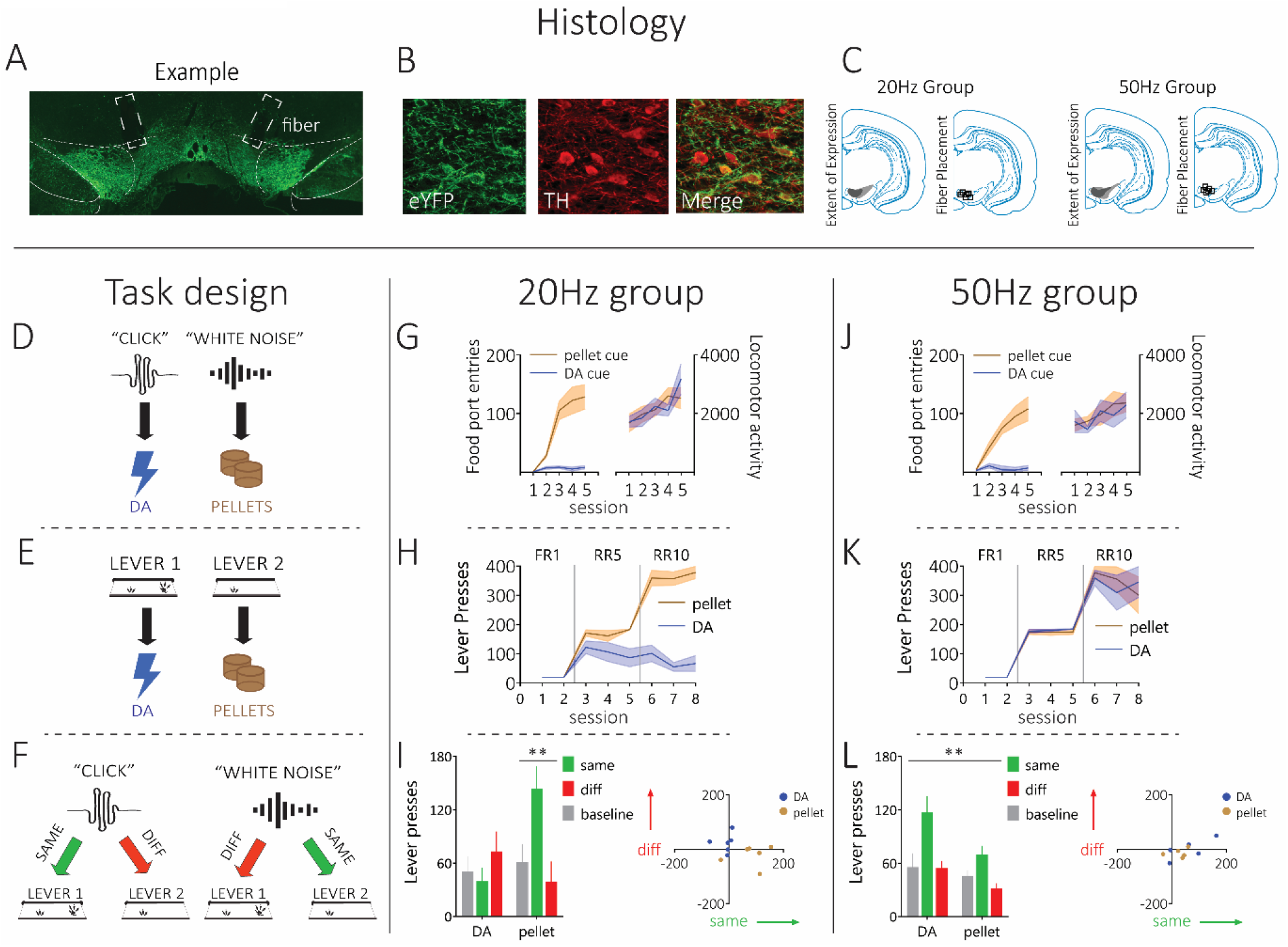
A physiologically-relevant frequency of dopamine stimulation (20Hz) does not function as a meaningful reward, however, high-frequency dopamine stimulation (50Hz) functions as a reward that is encoded as a specific sensory event. <u>Top</u>: Histological verification with **A**) bilateral Cre-dependent ChR2 expression in TH-Cre rats, **B**) colocalization of TH and virus expression approached ~90%, and **C**) schematic of minimum and maximum virus expression and fiber placement. <u>Left column</u>: Schematic illustrating the task design using one counterbalancing example, which consisted of **D**) Pavlovian conditioning, **E**) Instrumental Conditioning, and **F**) the PIT test. Rats first learnt that two auditory cues (e.g. click and white noise) led to two outcomes (e.g. dopamine stimulation and pellets), then they learnt to perform two lever presses that led to the two outcomes. Finally, rats were presented with the two auditory cues and given an opportunity to press either lever, without reward feedback. <u>Middle column</u>: **G**) Rats in the 20Hz group (*n*=6) showed an increase in food-port entries during the pellet-paired stimulus, but not the dopamine-paired stimulus. These rats showed equivalent increases in locomotor activity across learning to both stimuli. **H**) During instrumental conditioning, where rats learned to make lever presses for the two outcomes, rats in the 20Hz group showed robust lever-pressing responses for the pellets, but not the dopamine stimulation. **I**) In the final PIT test, when the pellet-paired cue is presented, these rats showed significant elevations in responding on the pellet-paired lever, indicating specific PIT. However, they did not show PIT for the dopamine-paired cue. <u>Right column</u>: **J**) Rats in the 50Hz group (*n*=5) showed increases in food port entries during the pellet-paired stimulus, but not the dopamine-paired stimulus. Increases in locomotor activity across learning was similar for both the dopamine- and pellet-paired stimulus. **K**) During instrumental training, the 50Hz group showed robust lever pressing for both the dopamine stimulation and the pellets. **L**) In the final PIT test, the dopamine- and pellet-paired stimuli both produced robust specific PIT. Error bars =SEM.

**Figure 2.**
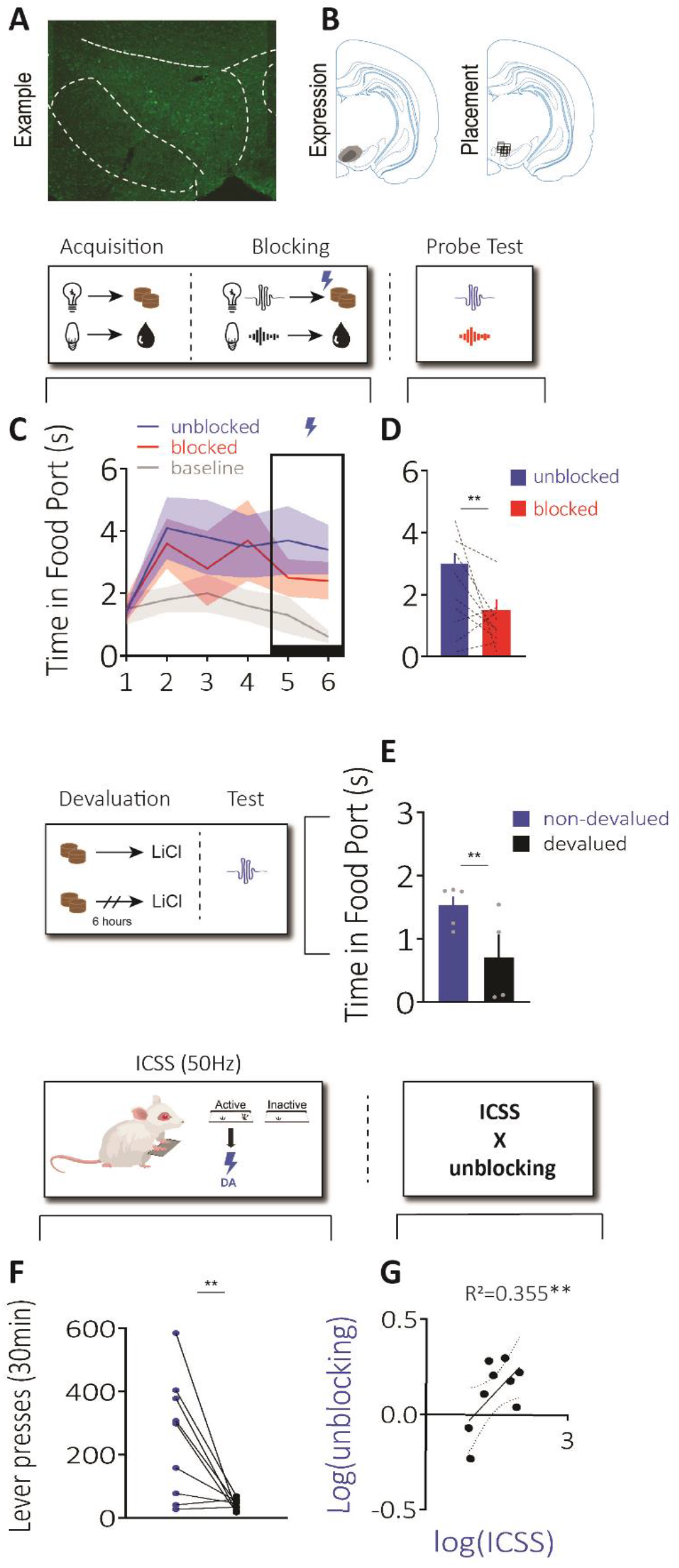
A physiologically-relevant frequency of dopamine stimulation (20Hz) functions as a teaching signal to drive sensory-specific learning. **A)** Unilateral example of bilateral ChR2 expression in dopamine neurons, **B)** Left: minimum and maximum expression of virus across rats, Right: placement of fiber tips across rats. <u>Top schematic</u>: design of our blocking task using one counterbalancing example, which consisted of acquisition, blocking, and a probe test. **C**) Rats (*n*=9) first learnt that two visual cues led to two distinct rewards (acquisition), then two novel auditory stimuli were introduced in compound with the visual cues and led to the same rewards (blocking). During blocking, we stimulated dopamine neurons (1s, 5ms, 20Hz) at reward delivery for one of the compounds (“unblocked” cue). Rats acquired the food-port response during the cues and maintained high levels of responding after introduction of the auditory cues and dopamine stimulation. **D**) We next gave rats a probe test and found they responded more to the unblocked cue, relative to blocked cue. <u>Middle schematic</u>: devaluation procedure used to devalue the reward paired with the unblocked cue. **E**) Rats showed a significant reduction in responding to the unblocked cue following devaluation, confirming dopamine stimulation unblocked the association between the cue and a sensory-specific representation of reward. <u>Bottom schematics</u>: design used for ICSS, rats were given access to a lever that produced 50Hz stimulation of dopamine neurons (active), or nothing (inactive). **F**) Rats pressed more on the active lever, **G**) This ICSS measure was positively correlated with the unblocking effect in B (r= 0.56; *p*=0.045). Error = SEM.

After this training, rats moved onto the instrumental contingencies where they could press one lever to receive dopamine stimulation, and another to receive sucrose pellets. During instrumental training, one lever was presented at a time and rats could earn a maximum of 40 rewards, in line with other PIT studies^59,60^. This also helped us to prevent competition between the two instrumental actions during learning. At first, it only took one lever press to earn these rewards (fixed ratio 1; FR1), then rats needed to increase the number of lever presses to an average of five (random ratio 10; RR5) and then ten (RR10) to earn the reward. This progressive schedule is typically used to establish a robust baseline of performance that will withstand the PIT test that takes place without reward deliveries^59,60^. During instrumental training, significant differences between our 20Hz and 50Hz groups emerged (Fig 1H and K). Specifically, while our 20Hz group reached criterion on FR1 schedules earning the stimulation, their performance was not robust when we moved to leaner schedules. This was despite their maintenance of responding on these schedules for the food reward. However, rats in the 50Hz group continued to show robust performance for both rewards as instrumental training progressed. This was confirmed with statistical analyses, where a repeated-measures ANOVA demonstrated a significant main effect of reward (dopamine vs. food: *F*_1,9_=21.528, *p*=0.001), and a reward x group interaction (*F*_1,9_=21.544, *p*=0.001), owed to a significant difference in responding for dopamine stimulation between the 20Hz and 50Hz groups (*F*_1,9_=30.738, *p*=0.000), which was not present when comparing responding for food rewards (*F*_1,9_=0.128, *p*=0.729). This same pattern was also seen when considering the escalation of responding across sessions, which returned a main effect of session (*F*_7,63_=149.418, *p*=0.000), a session by group interaction (*F*_7,63_=9.413, *p*=0.000), owed to a difference in responding emerging between groups as instrumental training progressed (e.g., session 8: *F*_1,9_= 7.629, *p*=0.022). There was also a between-group difference in overall levels of responding (*F*_1,9_=15.647, *p*=0.003). Thus, while 50Hz dopamine stimulation functioned to motivate vigorous responding that was comparable to food rewards, consistent with other reports^4,7,17,18,25,52^, 20Hz of dopamine stimulation did not support robust instrumental responding beyond a continuously-reinforced schedule and was not comparable to a food reward.

Following training, rats received the PIT test where the dopamine- and food-paired cues were played and rats had the opportunity to press either lever. During this test, neither the cues or the levers produced their associated reward, which allowed us to investigate the nature of the reward representation that was present during learning in the earlier stages of the task, in the absence of reward feedback^54,56^. In our 20Hz group, we found that the dopamine-paired cue did not produce specific PIT (Fig 1I). Specifically, rats did not elevate their responding on either lever when the dopamine-paired cue was presented, relative to baseline. However, when the food-paired cue was presented, these same rats selectively elevated responding on the lever that produced the food, demonstrating selective PIT. In contrast, in our 50Hz group, we found that the dopamine-paired cue supported robust selective PIT, which was comparable to that produced by the food-paired cue (Fig 1L). This was supported with statistical analyses. A repeated measures ANOVA comparing responding in our 20Hz group on the same and different levers, relative to the baseline period prior to cue presentation, revealed a cue by lever interaction (cue x lever: *F*_1,5_=15.383, *p*=0.011), owed to a selective increase on the same lever in response to the food-paired cue (same vs. diff: *F*_1,5_=13.957, *p*=0.013) that was not present for the dopamine cue (*F*_1,5_=3.759, *p*=0.110). Importantly, there was also a significant difference when comparing responding on the same lever for the dopamine-vs. the food-paired cue (*F*_1,5_=7.885, *p*=0.038), which was not seen for the different lever (*F*_1,5_=3.243, *p*=0.132). This same analysis conducted on responding exhibited by our 50Hz group revealed a main effect of lever (same vs. different: *F*_1,4_=20.193, *p*=0.011), but no main effect of cue (dopamine vs. food: *F*_1,4_=0.585, *p*=0.487), or any interaction between these factors (*F*_1,4_=0.505, *p*=0.516). Thus, stimulation of dopamine at physiologically-relevant levels (i.e., 20Hz) did not function as a specific reward that was capable of supporting instrumental lever pressing in the same way that a natural reward could, and did not endow the cue with motivational significance that could support PIT. However, stimulating dopamine at higher ranges (i.e., 50Hz), typically used for ICSS studies^4,52^, supported robust instrumental responding on leaner schedules and selective PIT. This demonstrated that stimulation of dopamine neurons at supraphysiological frequencies during ICSS produced a supraphysiological sensory event that was capable of acting as a representation of a specific reward to motivate behavior, over and above any role for this signal in endowing antecedent cues with general value.

Next, we wanted to confirm that our 20Hz stimulation of dopamine neurons was effective. To do this, we ran an additional experiment with a new cohort of rats to test whether our 20Hz stimulation of dopamine was capable of acting as a teaching signal to produce learning, as we and others have previously demonstrated^22,36,37,61^. Here, we used the blocking procedure. We first trained rats that two light cues led to two different rewards (Figure 2 “acquisition”; e.g. house light→sucrose pellet, flash→grain pellet; counterbalanced). Then, we presented the lights in compound with novel auditory cues, which produced the same rewards (e.g. Figure 2 “blocking”; house light + white noise → sucrose pellet, flash + click → grain pellets; counterbalanced). Usually, subjects do not associate the novel auditory cues with the rewards, because the rewards are already predicted by the lights, termed blocking^62^. However, we stimulated dopamine neurons (1s 20Hz) as one of the rewards was delivered. This created an “unblocked cue”, which was the cue paired with dopamine stimulation during its associated reward. We could then compare with our “blocked cue”, which was the cue that was not paired with stimulation of dopamine neurons. All rats learnt to enter the port while the visual cues were presented, increasing above baseline across learning, which was unaffected by introduction of the auditory cues or dopamine stimulation (Figure 2C; time period: *F*_2,16_=11.072, *p*=0.001, session: *F*_5,40_=1.389, *p*=0.249; time period x session: *F*_10,80_=2.710, *p*=0.006; unblocked vs. baseline: *p*=0.006; blocked vs. baseline: *p*=0.003, unblocked vs. blocked: *p*=0.140). This demonstrated that all rats learnt that the visual cues were predictive of reward, and that stimulating dopamine neurons did not interfere with the ability to continue to respond during the food-predictive cues.

We then tested responding to the auditory cues alone without reward to assess how much learning had accrued towards them (Figure 2 “probe test”). Here, we saw that rats spent more time in the food port during the unblocked cue relative to the blocked cue (Figure 2D; *F*_1,8_=3.614, *p*=0.047). This showed that stimulating dopamine neurons was capable of driving additional learning to the stimulation-paired unblocked cue. To further probe the nature of the association that had developed via dopamine stimulation, we devalued the reward paired with the unblocked cue (Figure 2 “devaluation”). To do this, we allowed all rats to consume the reward paired with the unblocked cue outside the experimental chamber. Then, we immediately injected half the rats with lithium chloride (LiCl) to induce gastric malaise (devalued group). In the other rats, we injected LiCl 6 hours after consumption of the reward (non-devalued group). This would allow us to test whether the learning produced by dopamine stimulation involved a sensory-specific representation of the reward. Specifically, we could then test responding to the unblocked cue again and see if it was impacted by devaluation. During this test, we found that rats in the devalued group spent less time in the food port during the cue (Figure 2E; group: *F*_2,7_=5.15, *p*=0.029). This was an effect not seen when we tested responding to the blocked cue, whose associated reward had not been devalued, confirming the sensory-specific nature of the effect (Fig S4, left; group: *F*_2,7_=0.157, *p*=0.352). Together, these data show that our physiologically-relevant stimulation parameters were capable of acting as a teaching signal to drive sensory-specific associations between events, despite not being able to function as a reward in itself.

In these same rats, we next wanted to ask whether the rewarding effects of high-frequency stimulation are related to the ability of these neurons to act as a teaching signal. To test this, we examined how much the rats would press to get optogenetic stimulation of dopamine neurons at 50Hz (Figure 3 “ICSS”). Unsurprisingly, we found that rats would press consistently for 50Hz of dopamine stimulation (Figure 3F: *F*_1,8_=10.42, *p*=0.006). We hypothesized that the degree of lever presses for 50Hz stimulation would be positively correlated to the unblocking effect (Fig 2D). Indeed, when we compared the magnitude of the unblocking with the degree to which high-frequency dopamine stimulation would support ICSS, we found that these factors were strongly positive correlated (Fig 2G: Pearson r = 0.596, R^2^=0.355, *p*=0.045). We did not see this same relationship with ICSS and the blocked cue (Fig S2, right: Pearson’s r=0.148, R^2^=0.022; p=0.352). These data support a conclusion that the differential effects of 20Hz and 50Hz stimulation parameters are due to the change in frequency of firing of these neurons, and not because these different frequencies are tapping into different populations of dopamine neurons.

**Figure 3.**
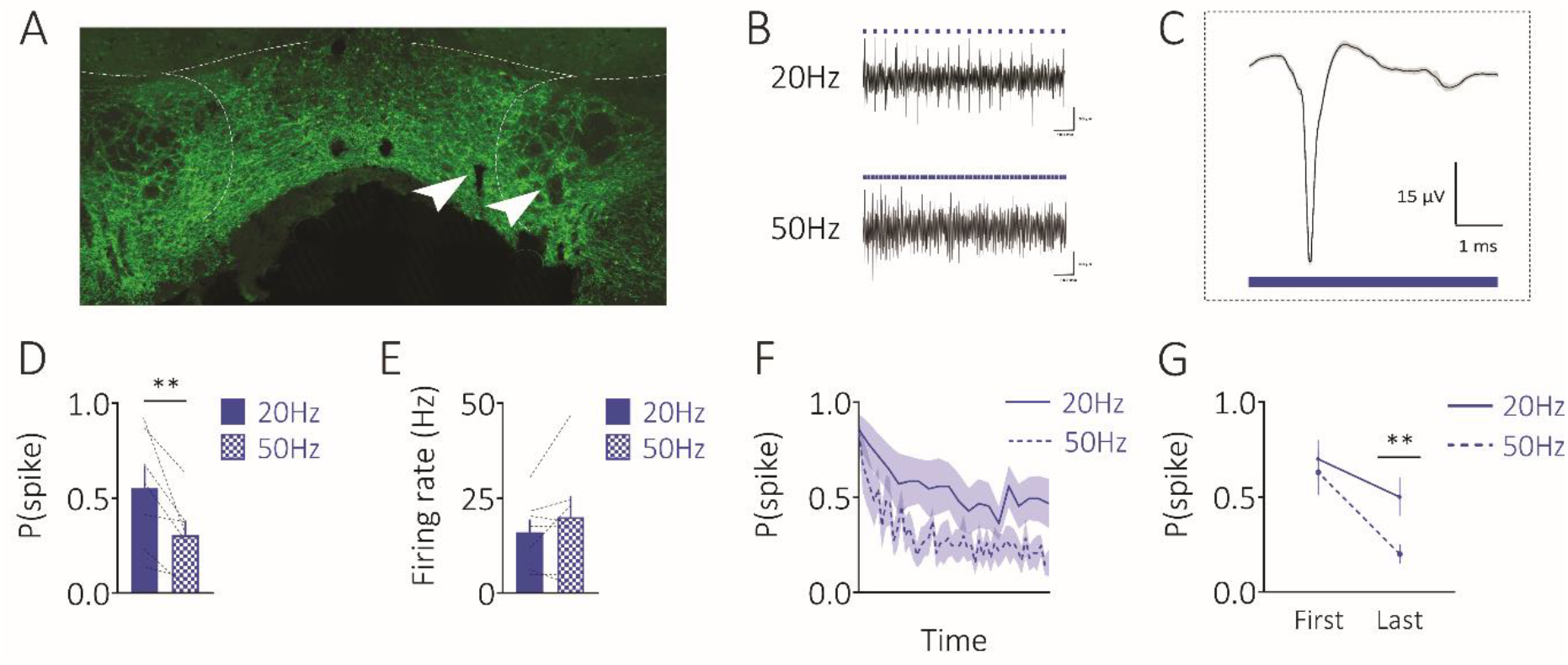
Optogenetic stimulation of VTA dopamine neurons at 20Hz and 50Hz results in the same number of action potentials across different timescales. Simultaneous recording and stimulation of VTA dopamine neurons reveal the firing dynamics produced by the two different stimulation trains. **A**) Example of bilateral Cre-dependent ChR2 expression (green) and placement of microwires, **B**) example trace of 20Hz and 50Hz stimulation trains, **C**) typical triphasic DA extracellular spike, **D**) neuronal activity follows the 20Hz stimulus train more faithfully then 50Hz, **E**) there is no difference in the overall number of action potentials generated by the 20Hz and 50Hz stimulus train, **F**) the rate of neuronal firing in VTA dopamine neurons decreases more rapidly across the 50Hz stimulation train relative to 20Hz, **G**) the difference in spike probability between the stimulus trains emerges towards the end of the stimulus train.

As a final experiment, we wanted to establish how VTA dopamine neurons were responding to our differential stimulation parameters. We infused a Cre-dependent ChR2 into the VTA of TH-rats. During this surgery, rats were implanted with a 16-channel microwire array with a central optic fiber into the VTA, which would afford simultaneous stimulation and recording of dopamine neurons. Similarly to prior results^17^, we found that the 20Hz stimulation produced firing rates of greater fidelity to the stimulation parameters than 50Hz (Figure 3A; *F*_1,6_=12.38, *p*=0.013). Interestingly, the firing rates of the neurons did not significantly differ overall across the 1s stimulus train (Figure 3B). However, there was a significant difference in the rate of decline in fidelity between the stimulation parameters across the stimulus train. There was no difference in the spike probability as a result of a pulse within the first 5 light pulses between the 20Hz and 50Hz stimulation parameters (Figure 3C-D; *F*_1,6_=4.7, *p*=0.074). However, a significant difference emerged towards the end of the stimulus train (Figure 3D; *F*_1,6_=11.61, *p*=0.014). These data show that both stimulation parameters produce a similar number of action potentials, but that the 50Hz stimulation parameters produce them across a shorter period of time. This demonstrates that it is the higher frequency, and not the greater number of spikes *per se* that generate the difference in the ability of these stimulation parameters to drive reinforcement.

Here, we investigated the psychological basis of ICSS with dopamine neurons. Somewhat surprisingly, there are very few studies that have asked this question, despite the high volume of manuscripts that have demonstrated ICSS throughout the brain. Using the PIT procedure, we asked whether stimulation of dopamine neurons functions to create a sensory-specific reward in its own right, much like a food reward, over and above the potential for this signal to endow the lever press or cue with general value. On this experimental backdrop, we also manipulated the frequency of stimulation between groups to see if a learning-relevant frequency of dopamine stimulation would function in the same way as high-frequency stimulation that is usually used to support ICSS. We found that 20Hz dopamine stimulation, relevant to physiological rates of phasic dopamine firing during learning^36,37,47,51^, would not support robust ICSS or serve as a sensory-specific event that would support the PIT effect. This was despite these same rats showing a significant selective PIT effect for a food reward. In contrast, 50Hz dopamine stimulation did support robust ICSS, even when the stimulation was made harder to receive on our leaner reinforcement schedules. The cue paired with 50Hz dopamine stimulation also supported selective PIT, where the dopamine-paired cue selectively increased responding on the lever that produced dopamine stimulation.

It is worth noting that we did not see any evidence for general value accumulating to the dopamine-paired cue or instrumental response in either our 20Hz or 50Hz group. That is, if dopamine did endow the cues or actions with general value, and particularly if this was the dominant mechanism, we would have expected the dopamine-paired cue to elevate responding on either lever above baseline, indicating the ability of this cue to generally invigorate instrumental responding in non-specific ways^58^. This is not what we saw. Instead, we saw in the 20Hz group that the dopamine-paired cue did not increase responding on either lever above baseline, and in our 50Hz group the dopamine-paired cue exclusively elevated lever-press responding on the same, congruent lever, and not on the different, incongruent lever. It is possible, however, that the presence of counterfactual information occluded any effect of general value in our PIT test^63^. Specifically, it has been demonstrated that Pavlovian training also produces inhibitory associations between the current cue that is being presented, and the outcome that is not currently being predicted by that cue (e.g., presentation of the dopamine-paired cue could signal the absence of the food), which reduces the likelihood that rats will increase responding on the incongruent lever. If this was the case, then general value could have been present, but this was overridden by the specific associations promoting the congruent action, and inhibiting the incongruent action. Given that we did not find evidence for specific PIT in our 20Hz group (i.e. there was no increase on the same lever or suppression on the different lever relative to baselines pressing rates), this is an unlikely reason that we did not see evidence of general value in this group. While it is possible that the specific associations occluded the ability to see general value accumulating to the cue and action in our 50Hz group, what is clear is that general value was not the dominant mechanism driving learning in this task and could not overshadow the specific associations. At the very least, these results suggest that the ability of supraphysiological dopamine stimulation to support ICSS does not necessarily support the value hypothesis of dopamine, and provide no evidence for this hypothesis at a physiologically-relevant frequency.

Importantly, we performed a control experiment to show that our stimulation was functioning effectively. Specifically, we showed that 20Hz of dopamine stimulation could produce learning, which we and others have shown previously^22,36,37,61^. In line with this, we found that 20Hz stimulation of dopamine was capable of producing learning about a sensory-specific cue-food associations when delivered as a prediction error during blocking. We showed this in both a novel cohort (Figure 2) as well as in the same rats that has received PIT (Figure S3). This demonstrated that our stimulation parameters were functioning effectively to produce a physiologically-relevant signal that could drive learning, and ineffective stimulation during PIT could not be a reason we found that 20Hz would not promote robust ICSS or the PIT effect. It is interesting to consider why 20Hz stimulation would produce any lever-press responding on a continuously-reinforced schedule, and also nose-poke responding in additional control experiments in line with other published findings^14^ (Fig S4), if it was not sufficient to support robust ICSS on leaner reinforcement schedules, or the PIT effect. This suggests that stimulation of dopamine neurons at 20Hz was reinforcing enough to promote some instrumental responding. Again, while we cannot rule out that this signal was generally valuable in some sense, what is clear is this value was not sufficient to support the kind of instrumental responding we see with other appetitive rewards, like food, water, or drugs of abuse. Indeed, we have explicitly demonstrated that a 20Hz signal delivered as a prediction error during learning will drive learning without making the antecedent cue valuable^36^, consistent with these data and explicitly against the formal predictions of the value hypothesis of dopamine as formalized by the model-free reinforcement-learning algorithm in temporal-difference reinforcement learning (TDRL).

These results are important for two reasons. Firstly, they show that frequency of dopamine stimulation relevant to reinforcement learning (i.e., 20Hz) does not function as a reward in and of itself in that it will not support robust ICSS or specific PIT. We also did not see evidence that the dopamine-paired cue generally invigorated responding on either lever during our PIT test, consistent with an inability of this signal to endow cues with general value, mirroring our previous work^36^. This is particularly pertinent to the field right now as we navigate new complex models to account for the seemingly discrepant function of dopamine neurons in different settings. These data now show that even in the context of ICSS, a prediction-error signal does not seem to function as a reward that possesses sufficient reinforcing properties to underlie that which is contained in other appetitive rewards, like food or drugs of abuse, necessary to uphold the value hypothesis. Secondly, these results also shed light on the psychological basis of ICSS using high frequencies of dopamine stimulation. Specifically, these data demonstrate that use of high-frequency dopamine stimulation in the context of ICSS functions to create a sensory event that acts as a specific reward in its own right. This does not have a basis in our everyday learning experience. Put simply, our learning experience does not contain representations of phasic dopamine as a rewarding sensory event. This questions the assumption that the ability of dopamine neurons to support ICSS at high frequencies indicates something about the region’s role in reinforcement learning. More generally, it begs the question: is there any physiological experience that might relate to ICSS? Perhaps a circumstance where this may become relevant is drug seeking^25,64^, where most drugs of abuse act in at least some way to increase phasic dopamine activity^65–67^. And if so, these data support an idea that people with substance use disorder seek out drugs of abuse to obtain a specific sensory experience, not because the actions or cues associated with the drug have become valuable.

## Author contributions

SM, KW, MJ, DB, and MJS designed the experiments. SM, IH, ZG, MHJ, and DJB conducted the experiments. SM and MJS wrote the paper with input from all authors. We thank Morgan Paladino for her assistance across the recording experiments.

## Data and materials availability

Data are available upon reasonable request.

## Supplementary Materials

Material and methods

Figs S1-S4

References

