## Supplemental materials for "The cognitive basis of intracranial self-stimulation of midbrain dopamine neurons"

**This PDF file includes:**

Materials and Methods
Figs S1-S4
References

### Materials and Methods

#### Subjects

Twenty-six experimentally naïve male and female Long-Evans transgenic rats carrying a Tyrosine hydroxylase (TH)-dependent Cre expressing system were used these experiments[1,2]. Rats were approximately 4 months of age prior to surgical procedures. Rats were maintained on a 12-hour light-dark cycle, where all behavioral experiments took place during the light cycle. Rats had *ad libitum* access to food and water unless undergoing the behavioral experiment during which they received sufficient chow to maintain them at ~85% of their free-feeding body weight. All experimental procedures were conducted in accordance with the Animal Research Council at UCLA and/or Animal Care and Use Committee at Rutgers University.

#### Surgical Procedures

Surgical procedures have been described elsewhere[1,2]. Briefly, rats were anesthetized with isoflurane (3%, 1-2% maintenance in 1L/m $O_2$) and secured in a stereotaxic device (David Kopf Instruments). Rats received two infusions of 1.0 µL bilaterally (2.0 µL per hemisphere) of AAV5-EF1α-DiO-ChR2-eYFP (n=24) into the VTA at the following co-ordinates: AP: -5.3mm; ML: ± 0.7mm; DV: -6.5 mm and -7.7 (females) or -7.0mm and -8.2mm (males). Virus was obtained from Addgene (MA, USA). For the behavioral experiments, optic fibers (Thorlabs, 200 µm diameter) were implanted bilaterally at the following coordinates relative to bregma: AP: -5.3mm; ML: ± 2.61mm and DV: -7.05mm (female) or -7.55mm (male) at an angle of 15º pointed toward the midline. For the recording experiment, a 16-channel microwire array with a central optic fiber was implanted targeting the VTA (-5 to -6.5 AP, +1.58 to 2.58 ML at 10º angle, -7.9 DV; Microprobes, Gaithersburg MD USA), per our previously published protocol[3].

#### Apparatus

Training was conducted in 8 standard behavioral chambers, which were individually housed in light- and sound-attenuating boxes (Med Associates, VT). For behavioral experiments, the chambers were equipped with a pellet dispenser that delivered 45-mg sucrose pellets (5TUT; BioServ, Frenchtown, NJ) into a recessed magazine when activated. Two retractable levers could be inserted into the chambers either side of the recessed magazine. Access to the magazine was detected by means of infrared detectors mounted across the opening of the recess. A computer equipped with medPC software (Med Associates, VT) controlled the equipment and recorded the responses. Raw data was output and processed in medPC2XL (Med Associates, VT) to extract relevant response measures. The chambers contained a speaker connected to white noise generator and a relay that delivered a 5 kHz clicker stimulus, as well as a house light that could illuminate the chambers when programmed.

#### Behavioral Procedures

##### *Pavlovian-to-Instrumental Transfer*

The Pavlovian-to-Instrumental transfer experimental procedure was based on that previously described[4,5]. Briefly, rats begin training with Pavlovian conditioning, which continued for 10 days. During the 60-min session, rats received four 2-min presentations of two cues (clicker or white noise, counterbalanced), each paired with one of two rewards (dopamine simulation or sucrose pellets, counterbalanced). Rewards were delivered at four points randomly throughout the 2-min cue. To stimulate dopamine neurons, light (473nm; 1s; 14-16mW) was delivered into the brain at either 20Hz or 50Hz (5ms pulses)[1,6]. Inter-trial intervals (ITIs) averaged 5 minutes in duration. Locomotor activity was recorded throughout the session and analyzed via eztrack[7]. Following Pavlovian conditioning, rats were given 8 days of instrumental training, where they were trained to lever press for pellets or dopamine stimulation. Each session consisted of two 10-minute blocks on each lever separated by a 2.5-minute time-out period in which levers were retracted. If an animal pressed a lever more than 20 times during a 10-minute session, then the lever was retracted immediately, and the 2.5-minute time-out period began before the next lever was made available, earning a maximum of 40 rewards on each lever. For the first two days of instrumental training lever presses were continually reinforced (CRF). Rats were then moved on to three days of a random ratio schedule (RR) where each levers delivered a reward with a probability of 0.2 (i.e. RR5), then finally to an RR10, which delivered a reward with a probability of 0.1. During the critical PIT test, both levers were extended, and no rewards were delivered. To extinguish instrumental responding on both levers, rats first received 8 minutes of extinction prior to cue presentation. Then, each cue was presented four times in the following order: clicker-noise-noise-clicker-noise-clicker-clicker, with a fixed 3-min ITI between cues. Rats received three such sessions, where they receive one RR10 session in between these tests.

*Blocking*

Rats first received presentation of two light cues followed by delivery of two distinct rewards (e.g. flash→sucrose; house light →pellet). Rats received 8 sessions, consisting of 14 trials, separated by variable ITI of 4mins. Then, rats received the visual cues in compound with two novel auditory cues to create two novel audiovisual compounds, which were followed by the same rewards (e.g. flash + click→ sucrose; house light + white noise → pellets). During compound training, rats received stimulation of dopamine neurons that coincided with reward delivery after one of the compounds[8,9] (473nm, 1s, 20Hz, 5ms pulses, 14-16mW), to mimic an endogenous prediction error[10-12]. Rats then received two probe tests with the click and white noise alone without reward, consisting of 8 trials with an ITI averaging around 4 minutes. All stimuli were counterbalanced as was the order of their presentation.

*Devaluation*

Following blocking, we trained rats on instrumental contingencies for the two distinct rewards. We then tested if the cues would promote PIT. However, we did not see significant PIT in any direction, primarily because rats spent a lot of time in the food port during the unblocked cue (blocked (± SEM): 1.5s (0.2); unblocked (±SEM): 2.7s (0.2); $F_{1,8}$=6.230, $p$=0.02). Instead, we conducted a devaluation test to assess the nature of the association that had developed between the unblocked cue and the reward. To do this, we allowed rats to consume the reward paired with the unblocked cue in a novel context for 30mins. Immediately after this, half the rats received injections of lithium chloride (LiCl; 0.15M, 10mg/ml; devalued group). The other half were returned to the colony room and received injections 6 hours later (non-devalued group). We repeated this for 3 days. After

48 hours recovery, rats were again placed in the experimental chambers and given a final probe test, where the unblocked cue was presented without reward. We also tested responding to the blocked cue (Fig S3). Each cue was presented for a total of 4 trials, with an ITI averaging around 4 mins.

*Intracranial self-stimulation*
To examine whether the rats from our blocking experiment would perform ICSS for 50Hz stimulation of dopamine neurons, we allowed them access to a lever that would produce 1s of 50Hz stimulation (473nm; 1s; 14-16mW, 5ms pulse) on a CRF schedule, and another than produced nothing. This session lasted for 30mins and rats could press the levers as much as they wanted.

**Recording Procedures**

Traditional photo-tagging procedures have accounted for differences in the amount of light reaching proximal and distal microwires by titrating the light delivery for each wire[13,14]. However, we opted against this approach, fixing the light intensity at 15mW, so as to provide an accurate model of neuronal firing in the tissue surrounding the optic fiber for the behavioral tasks used in the present paper. Accordingly, we observed a decrease in fidelity for neurons distal to the optic fiber, consistent with models of optogenetic stimulation. Neurons were classified as channelrhodopsin (ChR2)-responsive if they reliably exhibited responses within 15ms of the light onset[13]. Two confirm the absence of photoelectric artifacts, we recorded 2 additional rats with wires dorsal to the VTA. In these rats, we found no evidence optically evoked activity, even at optical power levels above those used in the experimental preparation.

On the day of testing, rats were places in a standard operant chamber and allowed to habituate for 10-15 minutes during which period neurons were identified and sorted online using Synapse software (TDT, Alachua, FL, USA). Single neurons were amplified and recorded using a digital headstage (ZD32, TDT, Alachua, FL, USA), preamplifier(PZ5), and bioamp (RZ2). Signals were then bandpass filtered at 300-5000Hz, and stored using Synapse (TDT, Alachua, FL, USA) at 24 kHz. Rats were first given 10 trials of 20 Hz stimulation (473 nm laser, 15mW, 5 ms pulse) with a 1-minute inter-trial interval, followed by 10 trials of 50 Hz stimulation (473 nm laser, 15mW, 5 ms pulse) with a 1-minute inter-trial interval.

**Histology**
*Behavioral experiments*
Rats were sacrificed by carbon dioxide and perfused with phosphate buffer followed by 4% paraformaldehyde in phosphate buffer. 20µm coronal sections were collected using a Cryostat (Leica Bio Systems), imaged and visualized for confirmation of virus expression and fiber tip placements using a Zeiss LSM 900 microscope with a 4x or 20x objective and ZEN Imaging software.

*Recording experiments*

Rats were terminally anesthetized and anodal current (50 µA, 4 sec) was passed through each microwire to mark the tip location. Rats were then perfused with phosphate buffer (PB, 0.1M, pH 7.4) followed by 4% paraformaldehyde (PFA). The brain was then stored in 4% PFA overnight before storing in 18% sucrose at 4 ºC until the brain equilibrated with the specific gravity of the sucrose solution. The VTA was then coronally sectioned at 30 µm to validate ChR2 expression and localize each microwire, as described previously[3]. Briefly, the positions of the microwires were identified by their tracks in the brain and their lesions at the microwire tips.

**Statistical Analyses**

Statistical analyses were conducted using SPSS 24 IBM statistics package. Analyses were conducted using a mixed-design repeated-measures analysis of variance (ANOVA). One-tailed tests were used for results with an a priori directional hypothesis. The data from the PIT test were analyzed across all trials of all three sessions of the PIT tests, and the blocking data were analyzed across the first six trials of both probe tests, consistent with other published demonstrations[8,15]. The ICSS and blocking data were correlated using Pearson's correlations on the log (x + 1) transformed data from the first 5 mins of the ICSS session and the ratio of responding to the blocked or unblocked cue relative to baseline (i.e. X - baseline/ X + baseline). For the recording studies, neuronal fidelity was calculated by examining the number of pulses within each trial where an action potential was recorded within 15 ms of the light onset. Fidelity was then further examined as a function of 1) each trial as a whole, 2) the pulse number in each train, and 3) the first 5 and last 5 pulses in each trial. The overall firing frequency across the 1 s train, irrespective of fidelity to individual pulses, was also analyzed. Data collection and analyses were not performed blind to the conditions of the experiments. Sample sizes were chosen based on similar prior experiments which have elicited significant results with a similar number of rats[16]. Further, we ran a formal post-hoc power analysis on the data elicited from these experiments to using G*Power 3.0 to estimate the power we had achieved using our sample sizes[17]. Specifically, we used the average of the partial $\eta^2$ (~0.8) from our analyses from the PIT test with the 20Hz and 50Hz groups to calculate the power (1-β) we had achieved. These analyses revealed an estimated power of 0.99 with the same sizes used in our study (Fig S1), with a type 1 error rate (α) below 0.05.

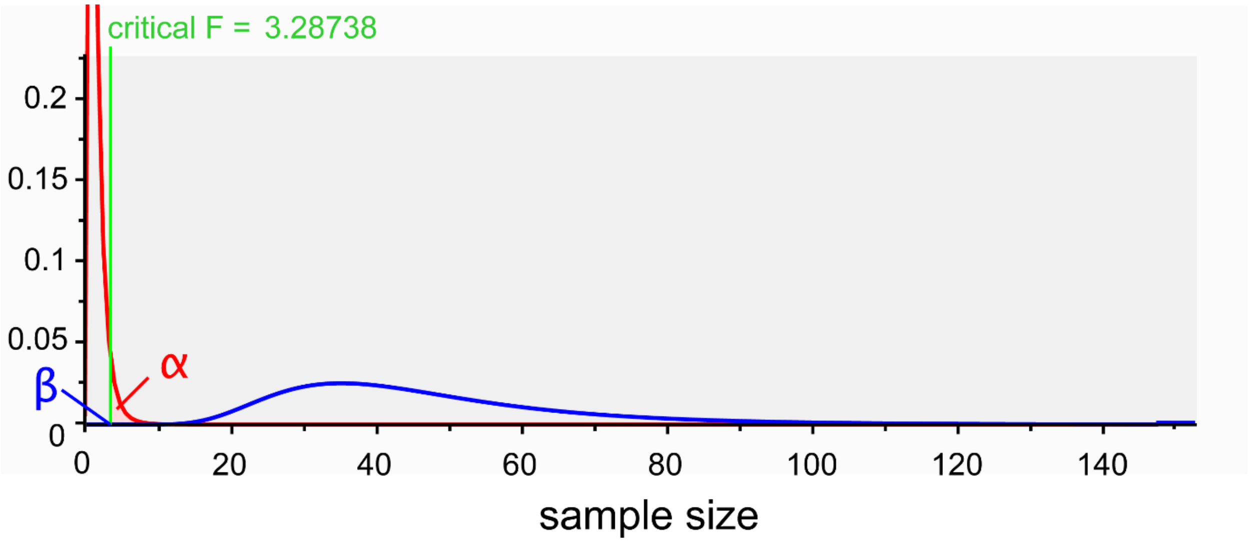


**Fig. S1. Formal power analyses showed that we obtained a high degree of power with the sample sizes used in these studies.** Using G*power 3.0[17], we conducted formal post-hoc power analyses on the data elicited from our PIT tests. The average partial $\eta^2$ elicited from our critical effects in our 20Hz and 50Hz groups was ~0.8, which resulted in a high degree of power (1-β; 0.99), and a type 1 error rate (α) below 0.05, with the sample sizes used in our study. This demonstrates that our sample sizes were sufficient to detect our critical effects with low likelihood of type 1 (α) or type 2 (β) errors.

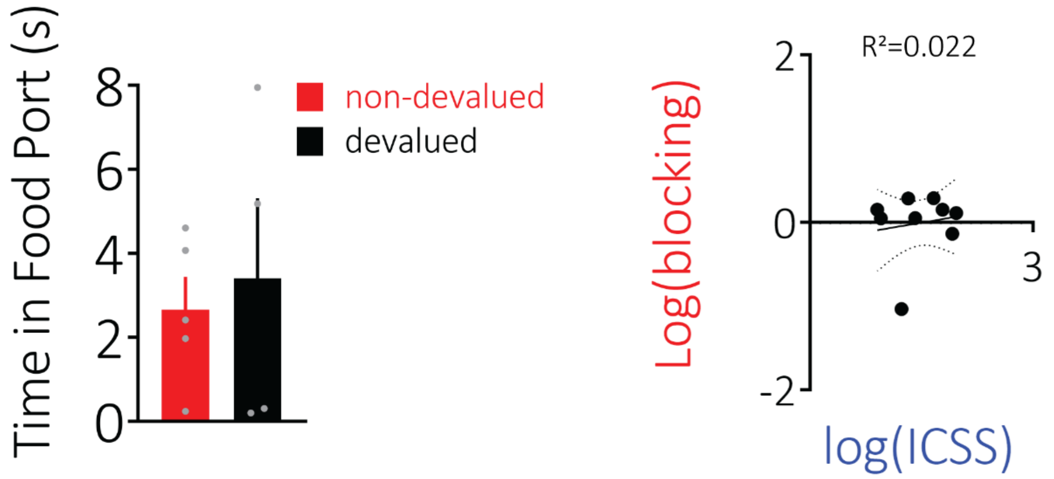


**Fig S2. Responding to the blocked cue is not sensitive to devaluation and is not correlated with ICSS.** Left: After devaluation of the reward paired with the unblocked cue, we tested responding to the blocked cue. We found that devaluation of the reward did not impact on responding to the blocked cue ($F_{1,7}$=0.157, $p$=0.352). Right: Responding to the blocked cue in the initial probe test illustrated in the main text in Figure 3B did not correlate with the degree to which rats would press the active lever to get stimulation of dopamine neurons (Pearson's r=0.148, $R^2$=0.022; $p$=0.352).

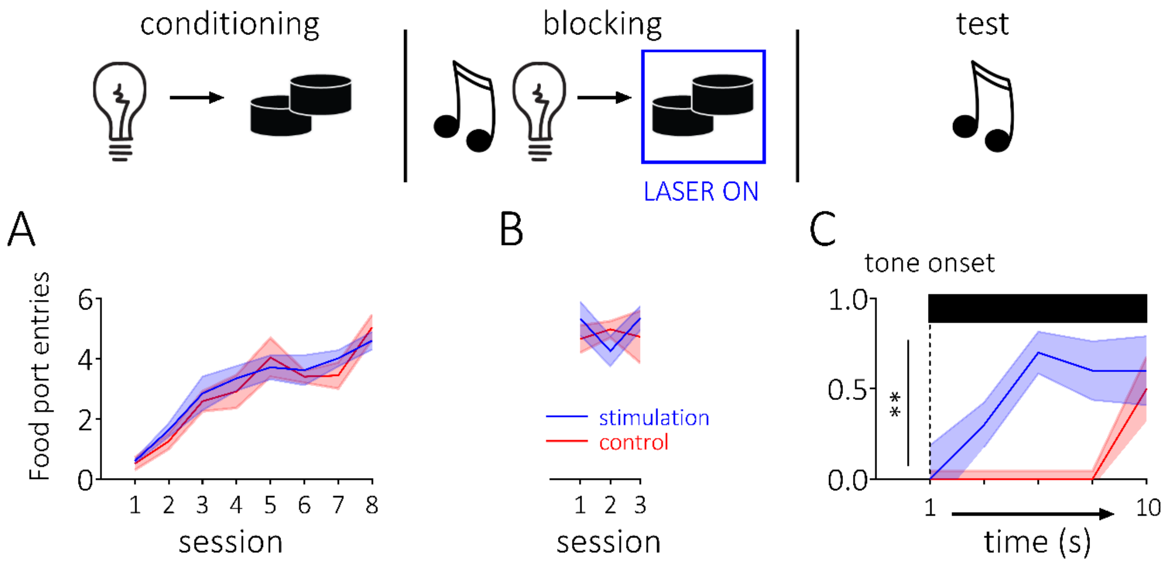


**Fig S3. A physiologically-relevant frequency of dopamine stimulation (20Hz) functions as a prediction error to unblock learning of cue-reward associations.** Top: schematic illustrating the task design. Bottom: food-port entries during conditioning (A), blocking (B), and the test (C). To ensure that our 20Hz stimulation was functioning to produce a physiologically-relevant signal, we tested whether these stimulation parameters would be sufficient to drive learning, as we and others have demonstrated previously. This experiment was conducted on the same rats that underwent the PIT experiment represented in Figure 2 of the main text. A) We trained rats (*n*=10) that a light led to delivery of two food pellets. All rats acquired the food-port entry response, with no difference between the groups (session: $F_{7,56}$=14.369, $p$<0.0001; session x group: $F_{7,56}$=0.344, $p$=0.930; group: $F_{1,8}$=0.119, $p$=0.739). B) Then, we presented a tone in compound with the light and the same food pellets. Usually, rats do not learn about the tone. However, when the food pellets were presented after the tone + light compound, we stimulated dopamine neurons at 20Hz in half the rats. All rats continued to exhibit high rates of responding at the food port, with no visible changes in these rates produced by dopamine stimulation (main effect, session: $F_{2,16}$=1.098, $p$=0.357; session x group interaction: $F_{2,16}$=1.608, $p$=0.218; group: $F_{1,8}$= 0.001, $p$=0.978). We then tested responding to the tone alone and found that dopamine stimulation had successfully unblocked learning (time: $F_{4,32}$=4.144, $p$=0.008; time x group: $F_{4,32}$=2.228, $p$=0.044; group (early): $F_{1,8}$=7.538, $p$=0.025), confirming our stimulation parameters were producing a physiologically-relevant dopamine signal. Error bars = SEM.

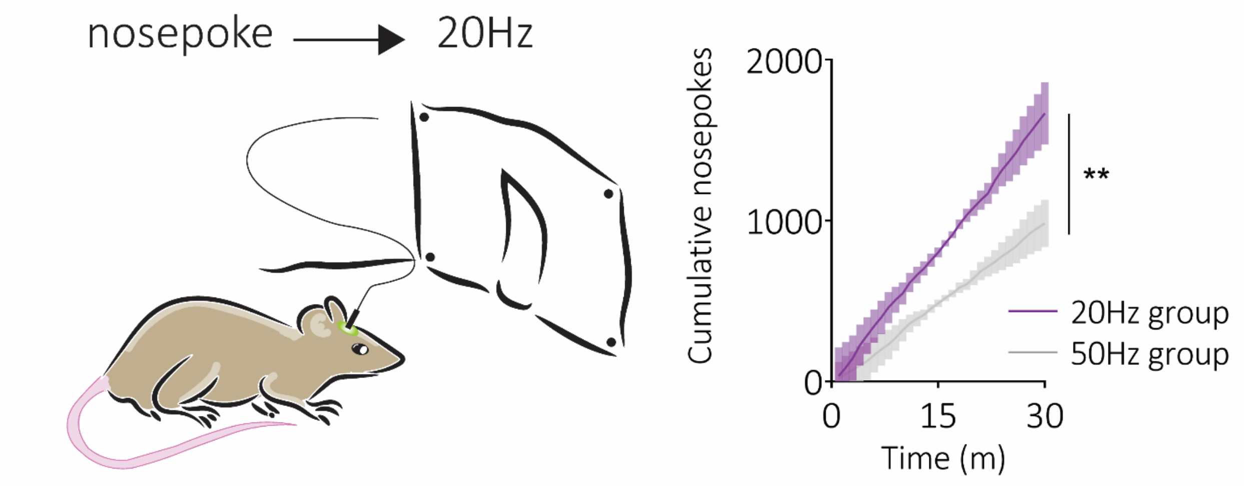


**Fig. S4. Rats will nosepoke robustly for 20Hz stimulation of dopamine neurons on a continuously-reinforced schedule.** We tested whether we could replicate the finding that rats will nosepoke at high rates for 20Hz dopamine stimulation[18]. Consistent with this data, we found that the rats that had previously received 20Hz of dopamine stimulation would nose poke robustly for 20Hz dopamine stimulation across the session. However, rats that had previously received 50Hz of dopamine stimulation would nose poke for 20Hz, but significantly less than the 20Hz group. This suggested that reducing the frequency of dopamine stimulation from 50 to 20Hz devalued the stimulation. This was confirmed with statistical analyses on the cumulative data, which showed a main effect of time ($F_{29,232}$=30.830, $p$=0.000), and an interaction by group ($F_{29,232}$=1.798, $p$=0.010), demonstrating that our 20Hz group increased responding across time at a greater rate than the 50Hz group.
